# PHASOR: An Open-Source Pipeline for Hidden Markov Model Analysis of Stimulus-Locked Retinal Multielectrode-Array Population Activity

**DOI:** 10.64898/2026.10.27.762251

**Authors:** Mohammed Junaid Shaik, Sichen Chen, Jessie Jin, Darwin Babino

## Abstract

Multielectrode-array recordings of the retina reveal neural population activity at cellular resolution, but relating recurring population states to stimulus timing and single-unit responses requires several linked analysis steps. We present PHASOR (Population HMM Analysis of Stimulus-locked Output in the Retina), a pipeline that combines stimulus-clock alignment, Fourier-based response classification, hidden Markov modeling with Chow-Liu tree emissions, model-order selection, and state visualization. We evaluated the workflow with synthetic populations and demonstrated its use in ten ex vivo recordings from three wild-type retinas (two male, one female) and one male rd1 retina, including 2 Hz full-field sinusoidal stimulation. Across ten synthetic datasets containing seven generative states, selected model orders ranged from seven to nine, with a mean of 7.9, matching the generative count exactly in three of the ten datasets. A representative confusion matrix showed a near one-to-one correspondence between generative and decoded states, with a single additional low-occupancy state, illustrating the distinction between recovering temporal structure and recovering the exact state count. In a representative retinal recording, response classes differed in stimulus phase and contrast dependence, and decoded states differed in occupancy, dwell time, and active-unit composition. Recordings during L-AP4 application, without projected stimulation, and in a photoreceptor-degenerate *rd1* retina illustrated changes in the resulting population-state descriptions. PHASOR integrates established analysis methods into a reusable workflow for relating stimulus-associated population activity to functional response classes. The selected states provide a descriptive model of population activity; their number and correspondence to biological assemblies require separate validation.

**Significance Statement:** Neurons respond together, yet analyses of individual cells can miss patterns in their collective activity. PHASOR connects recordings of many retinal neurons to a compact description of recurring population states and the response classes active within them. It brings stimulus alignment, response classification, model fitting, and visualization into one workflow. Synthetic examples show why a useful population description need not recover the exact number of states that generated the data. Retinal examples illustrate how the outputs relate to stimulus contrast and pharmacological perturbation. The tool is intended to make these analyses easier to repeat and inspect, while keeping model-derived states distinct from independently established biological cell types or assemblies.

## Introduction

Multielectrode arrays (MEAs) record the spiking activity of many neurons simultaneously, permitting study of how populations, rather than individual cells alone, represent a stimulus. In the retina, MEAs capture the output of retinal ganglion cells (RGCs), the neurons that transmit visual information to the brain. Collective activity carries information about visual input beyond that available from single cells (Warland et al., 1997) and can be organized into recurring, error-robust modes (Prentice et al., 2016; Loback et al., 2017).

Hidden Markov models (HMMs) provide a principled framework for describing such recurring patterns (Baum and Petrie, 1966; Rabiner, 1989). An HMM assumes that, at each time point, the population occupies one of a few hidden states, each characterized by a distinct profile of units likely to fire or remain silent. The model estimates the probability of remaining in a state or transitioning to another. Once fitted, it assigns the most probable state to each time bin, reducing the activity of hundreds of units to a compact state sequence.

Interpreting a state requires knowing which cells contribute to it. RGCs are functionally diverse, and the classification of an individual cell depends on the stimulus and response features examined (Baden et al., 2016; Jouty et al., 2018). Periodic stimulation provides a controlled means of characterizing this diversity, because each unit’s cycle-averaged response can be summarized by its polarity (ON or OFF), kinetics (transient or sustained), and phase relative to the stimulus (Chichilnisky, 2001; Gollisch and Meister, 2010). These response classes are treated here as descriptive labels rather than independently established cell types; assigning them to units allows each state to be interpreted through its constituent cells.

Applying HMMs to stimulus-locked recordings requires several linked steps. Each state requires an emission model specifying which binary units tend to fire together; Chow-Liu trees offer a sparse approximation based on the strongest pairwise dependencies (Chow and Liu, 1968; Schneidman et al., 2006). The number of states must also be selected, a step complicated by the tendency of predictive likelihood to favor more states than the generative process requires (Pohle et al., 2017; Li and La Camera, 2025). Finally, states must be related to unit responses.

Existing resources address parts of this workflow: PRANAS (Cessac et al., 2017) and Pyret (Naecker et al., 2017) support retinal spike-train and receptive-field analysis, HMM-spikes (Li and La Camera, 2025) fits Poisson HMMs, SpikeInterface (Buccino et al., 2020) handles spike sorting, and retinal population-mode analyses (Prentice et al., 2016; Loback et al., 2017) established tree-structured HMMs. PHASOR assembles stimulus synchronization, Fourier-based response classification, Chow-Liu-tree HMM fitting, model-order selection, and reporting into a single, documented, reusable workflow. PHASOR introduces no new statistical model; its contribution is integration.

We use synthetic data with known ground truth to evaluate state selection and generative-state recovery, then demonstrate the outputs in retinal recordings during 2 Hz sinusoidal stimulation across contrast levels, during pharmacological perturbation with L-AP4, and in two controls: no stimulus projected onto an intact retina, and full stimulus onto an *rd1* retina.

## Materials and Methods

### Software overview

PHASOR is organized into modules that mirror the phases of the analysis (Fig. 1): pre-processing, which aligns spikes to the stimulus clock and assembles the four-dimensional response array; processing, which partitions the data by stimulus condition and runs unsupervised uniform manifold approximation and projection (UMAP; McInnes et al., 2018) followed by OPTICS (ordering points to identify the clustering structure; Ankerst et al., 1999) clustering; Fourier classification, which assigns each unit its stimulus-locked polarity and kinetic label and records that unit’s cluster assignment alongside its label; HMM fitting, which searches hyperparameters, fits the model, and decodes latent states; and reporting, which renders the decoded states as a set of standardized figures. Default Fourier classifier parameters are listed in Table 1. The analysis backend is implemented in Python: each stage is an independent command-line program that reads and writes files in the pipeline’s documented on-disk formats, so that later stages, or an independent analysis, can consume its output directly, and a stage that has already produced a valid result, such as a fitted model bundle, is skipped rather than re-run.

**Table 1.** Default parameters of the cell-level Fourier polarity and kinetic classifier, with the role each parameter plays in the classification procedure described in Materials and Methods.

| Parameter | Value | Role |
|---|---|---|
| θ<sub>ρ</sub>: biphasic-ratio threshold | 0.6 | Biphasic vs. ON/OFF branch split |
| a<sub>min</sub>: per-cycle amplitude floor | 0.01 | Harmonic must be present in every<br>analyzed cycle |
| θ<sub>jit</sub>: phase-jitter threshold | 35° | Maximum cycle-to-cycle phase variability |
| θ<sub>SNR</sub>: SNR threshold | 0.3 | Minimum harmonic-reconstruction<br>signal-to-noise ratio |
| W<sub>thr</sub>: width threshold | 95 ms | Transient vs. sustained split |
| ε: numerical-stability constant (classifier) | 1×10<sup>−9</sup> | Added to denominators in the biphasic-ratio (A2/A1), SNR, and effective-width<br>calculations to avoid division by zero |
| peak_floor: minimum peak z-score | 0.5 | Gate on effective-width/kinetic-label<br>computation |

**Figure 1.**
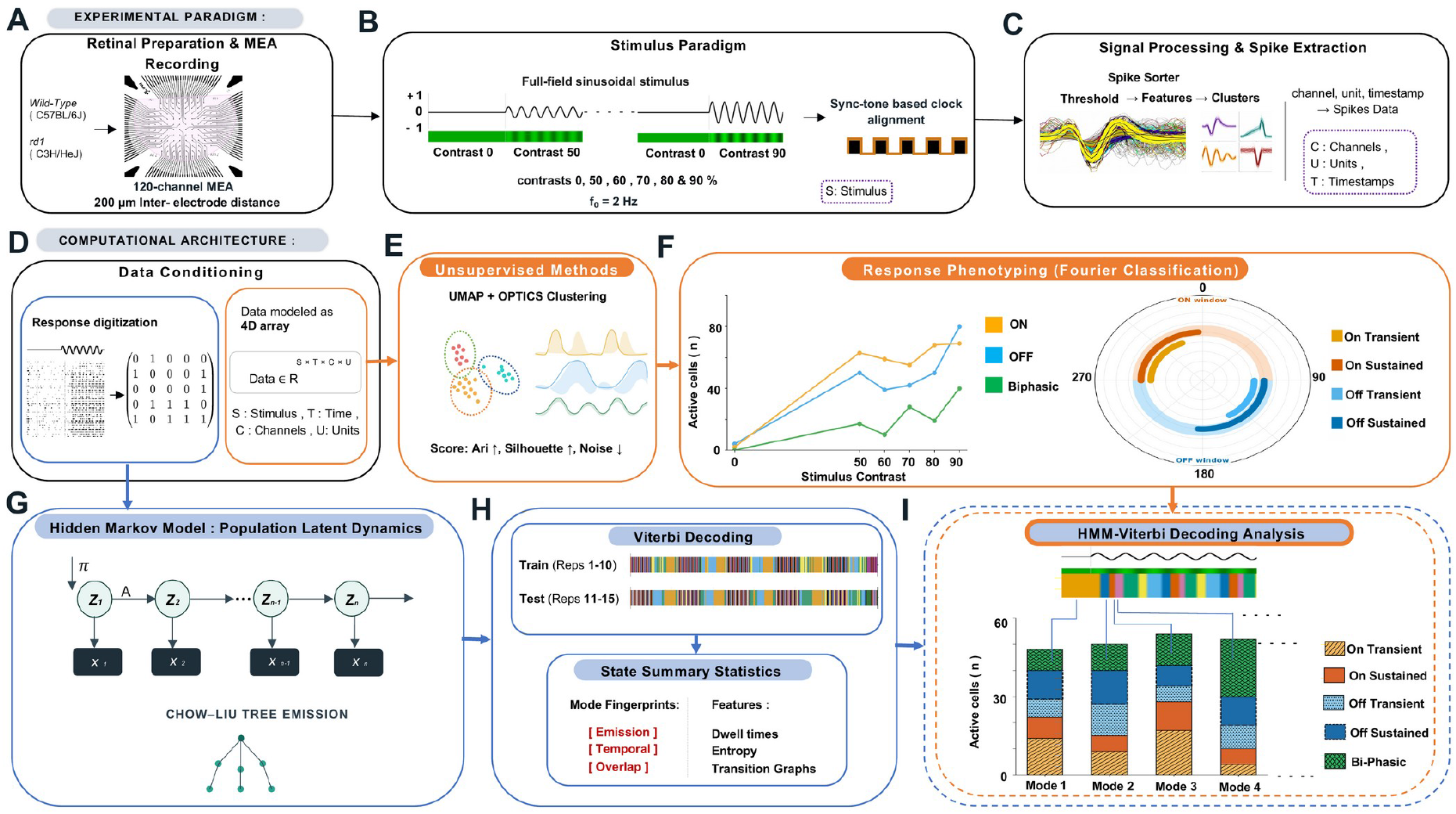
Overview of the PHASOR pipeline for population hidden Markov model (HMM) analysis of retinal ganglion cell activity, from data acquisition to state-resolved single-cell interpretation. (A) Ex vivo multielectrode array (MEA) recordings from wild-type (C57BL/6J) and rd1 (C3H/HeJ) mouse retinas using a 120-channel array with 200 µm inter-electrode spacing. (B) Full-field sinusoidal contrast stimuli (S) (f0 = 2 Hz) presented at six contrast levels (0, 50, 60, 70, 80, 90%). A synchronization signal, delivered on a dedicated stimulus channel, aligns the stimulus timeline with the MEA recording clock. (C) Spike sorting: threshold-based event detection, feature extraction, and unsupervised clustering yield a channel/unit/timestamp spike table for each identified single unit. (D) Data conditioning marks the transition from the experimental paradigm (A-C) to the computational architecture (D-I). Spike data are conditioned into two representations: a digitized response matrix used for HMM fitting and decoding (G-I), and a four-dimensional response tensor, Data ∈ ℝ^(S×T×C×U) (S, stimulus; T, time; C, channel; U, unit), used for single-cell response typing (E, F). (E) Unsupervised response typing: UMAP embedding followed by OPTICS clustering, evaluated by adjusted Rand index, silhouette score, and noise-point fraction. (F) Fourier-based response classification, independent of the clustering in (E). Left: active cell counts (n) as a function of stimulus contrast for ON, OFF, and biphasic classification using Fourier transform. Right: polar distribution of response phase across the ON and OFF stimulus windows, partitioning units into On-transient, On-sustained, Off-transient, and Off-sustained subtypes (polarity follows phase; transient/sustained labels use the response-width criterion in Materials and Methods). (G) Hidden Markov model: population activity xt at each time bin is generated by a latent state sequence z1,…,zn governed by initial distribution π and transition matrix A. The emission distribution P(xt|zt) is factorized over a Chow-Liu tree fit to pairwise mutual information between units. (H) The globally optimal latent state sequence is decoded via the Viterbi algorithm, applied separately to training (repetitions 1-10) and held-out test (repetitions 11-15) trials. Each decoded state is summarized by emission, temporal, and overlap fingerprints, together with dwell time, entropy, and transition-graph features. (I) Response-class composition of decoded states: Viterbi-assigned state membership (H) is combined with Fourier-derived response subtype (F) to determine, for each state, the number of active cells belonging to each subtype: On-transient, On-sustained, Off-transient, Off-sustained, and biphasic.

A browser-based interface, built with Next.js, presents the pipeline as a connected flow diagram organized by phase, with an in-page search for jumping directly to any stage. Each stage’s page lets the user locate its required input files through a live-filtered path picker and launch the stage, streaming its console output back to the browser as it executes. The HMM stage has a dedicated page that discovers the appropriate fitting workflow, launches the hyperparameter search, refit, and Viterbi decoding as a single detached background process, and reports progress through those three steps, resuming from whichever searches are already completed if the process is interrupted. Because the interface runs on the user’s own machine and issues the same command-line invocation a user would type directly, browser- and terminal-launched runs execute the same programs with the same arguments.

### Experimental data acquisition

Recordings analyzed with PHASOR in this study were obtained from adult C57BL/6J (wild-type) and C3H/HeJ (*Pde6b*^*rd1*^*/Pde6b*^*rd1*^, *rd1*) mice of both sexes, obtained from The Jackson Laboratory (Bar Harbor, ME). All animal procedures were performed in accordance with the institutional animal care committee’s regulations. Animals were maintained on a 14 h light/10 h dark cycle (lights on, 6:30 A.M.-8:30 P.M.) with food and water available ad libitum, group housed (2-4 per cage), and dark-adapted for at least 1 h before experiments, which were performed under dim red light during ZT14-ZT24 (ZT0, lights on). Mice were killed with CO_2_ and their eyes enucleated and placed in cold artificial cerebrospinal fluid (ACSF: 125 mM NaCl, 2.5 mM KCl, 1.2 mM NaH_2_PO_4_, 1 mM MgCl_2_, 2 mM CaCl_2_, 26 mM NaHCO_3_, 20 mM D-glucose, bubbled with 95% O_2_/5% CO_2_). Retinas were isolated, and the ventral segment was mounted ganglion-cell-layer-down on a 120-channel multielectrode array (120MEA200/30iR-Ti-gr; Multi Channel Systems, Reutlingen, Germany), kept at room temperature, continuously perfused with ACSF at 3.2 mL/min, and allowed to settle and recover for at least 1 h before recording. A subset of recordings included pharmacological perturbation with L-AP4 (2.5 µM; Tocris Biosciences, Bristol, UK; cat. 0103), an mGluR6 agonist that suppresses ON bipolar cell signaling (Slaughter and Miller, 1981; Nakajima et al., 1993), applied for 30 min before recording and followed by a 1 h washout. A separate subset of recordings was collected with no visual stimulus projected onto the retina (no-projection control; see Results). Four animals contributed the recordings analyzed with PHASOR in this study, one retina per animal: three wild-type (C57BL/6J) retinas (2 male, 1 female; age P125-P199) and one *rd1* (C3H/HeJ) retina (male; age P105), for 10 MEA recordings in total. These four retinas are a subset of a larger collection of 11 retinas; none of the ten recordings was excluded. WT1 (male, right retina, P199) and WT2 (female, left retina, P132) each provided four sequential recordings (pre-treatment run 1, pre-treatment run 2, 2.5 µM L-AP4, and washout) used for the pharmacological demonstration. WT3 (male, right retina, P125) provided a single no-projection control recording. The *rd1* retina, *rd1*-1 (male, left retina, P105), provided a single recording under the standard stimulus protocol and is included as a comparison recording. All ten recordings, with their animal metadata and fitted model hyperparameters, are listed in Table 2. Visual stimuli were delivered from below the array (presented onto the retina from the ganglion-cell side, following the direction of light entry in vivo) with a Digital Light Projector (LightCrafter 4500, Texas Instruments, Dallas, TX) through a custom optical lens system, providing 1280 × 800 pixels of spatiotemporally patterned stimulus over the full MEA area at a 60 Hz refresh rate; only the green LED channel (510 nm) was used, at an intensity of 2670 photons/µm_2_/s. Stimuli were authored and exported as synchronized video files using the pipeline’s companion stimulus-generation module. The primary stimulus was a 2 Hz full-field sinusoidal contrast flicker presented at 0, 50, 60, 70, 80, and 90% contrast, and a periodic synchronization tone was embedded in the recording to align the clocks of the stimulus computer and the MEA acquisition system. Each contrast level was presented as a 3000 ms trial (six 500 ms cycles) immediately preceded by a 3000 ms trial at 0% contrast; trials were grouped into four blocks of consecutive repetitions separated by 120 s of rest, following a 10 s adaptation period at the start of the protocol; this is the same stimulus block used to generate the synthetic validation data (see Simulated validation data below). Raw voltage traces were high-pass filtered (Butterworth, 350 Hz cutoff) and digitized at 10 kHz; spikes were detected at an amplitude threshold of -3.50 SD from the peak-height histogram distribution and automatically sorted using established spike-sorting methods (Lewicki, 1998) in a two-dimensional feature space with the Standard Beta Means algorithm (β = 1; Offline Sorter V4, Plexon, Dallas, TX), yielding, for each recording, a list of channel, unit, and spike time, the raw input to PHASOR’s preprocessing stage below.

**Table 2.** Recordings analyzed with PHASOR. One row per MEA recording (10 recordings from 4 animals, one retina per animal), giving animal label, sex, retina side, age at the time of the experiment, recording condition, the number of sorted single units entering the HMM, and the fitted hyperparameters: selected state count K, covariance soft-threshold η, and pseudocount α. η was selected per recording by cross-validation over the grid [0.0005, 0.002, 0.005]; α was held fixed at 0.5. WT1-WT3 are C57BL/6J and rd1-1 is C3H/HeJ.

| Recording | Sex | Retina | Age (P) | Condition | Units (N) | K | η | α |
|---|---|---|---|---|---|---|---|---|
| WT1 run 1 | M | Right | 199 | Pre-treatment | 276 | 12 | 0.005 | 0.5 |
| WT1 run 2 | M | Right | 199 | Pre-treatment | 288 | 12 | 0.005 | 0.5 |
| WT1 L-AP4 | M | Right | 199 | L-AP4, 2.5 µM | 318 | 12 | 0.005 | 0.5 |
| WT1 washout | M | Right | 199 | Washout | 274 | 12 | 0.005 | 0.5 |
| WT2 run 1 | F | Left | 132 | Pre-treatment | 259 | 12 | 0.005 | 0.5 |
| WT2 run 2 | F | Left | 132 | Pre-treatment | 257 | 15 | 0.005 | 0.5 |
| WT2 L-AP4 | F | Left | 132 | L-AP4, 2.5 µM | 256 | 12 | 0.005 | 0.5 |
| WT2 washout | F | Left | 132 | Washout | 248 | 12 | 0.005 | 0.5 |
| WT3 | M | Right | 125 | No projection | 260 | 12 | 0.005 | 0.5 |
| rd1-1 | M | Left | 105 | Standard stimulus | 255 | 12 | 0.005 | 0.5 |

### Preprocessing

PHASOR expects, as input, spike-sorted timestamps (channel, unit, spike time), a stimulus protocol file recording the timing and identity of every stimulus presentation, and a periodic synchronization (“synctone”) signal recorded on an auxiliary channel of the same MEA system, at intervals corresponding to stimulus events. Because the stimulus computer and the MEA acquisition system run on independent clocks, the raw recording is first scanned for every synctone pulse; each stimulus presentation is then anchored to its detected pulse and spike times are expressed relative to that pulse, so clock drift cannot accumulate across the recording (no separate drift-correction fit is applied). The pulse-anchored spike times, together with the parsed stimulus protocol, are then binned into fixed time windows and stacked into a four-dimensional array indexed by [Stimulus × time bins × channels × units]; for a periodic stimulus at frequency f0, the bin width and cycle length are chosen so that an integer number of bins spans one stimulus cycle (10 ms bins and 50 bins/cycle for f0 = 2 Hz in the recordings analyzed here). This four-dimensional array is consumed by the clustering and Fourier classification stages; the HMM stage builds its binary matrix from the same pulse-anchored spike times (see below). As a quality-control step, the pulse detector prints the mean, min, max, and SD of the deviation of each inter-synctone interval from its nominal 1 s spacing (in ms), and recordings with excessive timing error are flagged by visual inspection; these values do not alter spike times, and there is no automated numeric cutoff coded into the pipeline. Typical mean timing error is approximately 1.5 ms.

As an independent, per-recording quality-control check on cluster stability, the processing stage partitions the four-dimensional response array by stimulus condition and applies unsupervised UMAP (McInnes et al., 2018) followed by OPTICS (Ankerst et al., 1999) clustering. Each sample is one unit’s trial-averaged, smoothed, z-scored 300-bin response (see Fourier classification below). UMAP is run with a minimum distance of 0.05 and a correlation-based distance metric, and OPTICS with Euclidean distance. Tuning proceeds in two stages: stage 1 searches UMAP dimensionality (3, 5, 8) and number of neighbors (5, 10, 15) with default OPTICS settings (minimum samples for a core point = max(5, 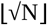), minimum cluster size = max(5, ⌊0.05N⌋), steepness 0.05, where N is the sample count); stage 2 searches OPTICS minimum samples (30, 50, 70), minimum cluster size as a fraction of N (0.003, 0.005, 0.01; at least 5), and steepness (0.03, 0.05, 0.08) at the selected UMAP setting. Every combination is run with seeds 0-4, and the final embedding uses seed 42. Each parameter combination is evaluated by the adjusted Rand index (ARI), silhouette score, and noise-point fraction. ARI is computed pairwise across random seeds, so it measures clustering stability rather than agreement with a ground truth; silhouette score and noise fraction are medians across seeds. A combination is accepted if median ARI > 0.85, median silhouette > 0.45, and noise fraction < 0.45, and among accepted combinations the one with the lowest noise fraction is selected, then the highest silhouette, then the highest ARI. If no combination is accepted, the best-ranked non-degenerate combination (at least two clusters, positive silhouette, nonzero noise fraction) is used and reported as not meeting the criteria.

For HMM fitting, spike times are discretized using the same *Δt* = 10 ms bins defined above, and, for each neuron and bin, activity is represented as a binary variable indicating whether the neuron fired at least one spike in that bin, giving a binary population activity matrix *X* ∈ {0,1}^*T×N*^, where *T* is the number of time bins and *N* is the number of recorded units.

### Fourier classification of stimulus-locked cell subtypes

To interpret decoded HMM states in terms of the functional response classes that contribute to them, every recorded unit is independently assigned a stimulus-locked functional label from a Fourier analysis of its own cycle-averaged response, using only the harmonics fixed by the stimulus frequency, following the general logic of frequency-domain characterization of light response (Chichilnisky, 2001; Gollisch and Meister, 2010). Labels are computed separately for each contrast condition from the trial average over all 15 repetitions, including the held-out repetitions; they are used only to interpret decoded states and do not enter HMM fitting.

For each unit, spike counts are trial-averaged into a peri-stimulus time histogram, smoothed in time with a Gaussian kernel (σ = 3 time bins), and z-scored per unit using the mean and SD (a separate numerical-stability constant, 1e-8, added to the SD; distinct from the classifier’s ε in Table 1, which stabilizes a different set of denominators) so that classification depends on response shape rather than firing-rate magnitude. Units with no spikes in a condition are omitted, and a matched-size 0% contrast baseline is subtracted before classification where available. (Smoothing and z-scoring are performed in an upstream preprocessing stage that produces the per-unit response array consumed by the classification stage described below.) Folding the resulting trace into P-bin cycles (one stimulus cycle), discarding the first cycle to avoid the onset transient, and averaging the remaining C cycles gives the cycle-averaged response:

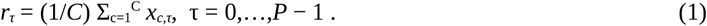

Here, *x*_*c,τ*_ is the z-scored, cycle-folded response at cycle *c* and within-cycle bin *τ* (i.e., the smoothed, trial-averaged peri-stimulus histogram described above, after folding into *P*-bin cycles).

Its discrete Fourier transform,

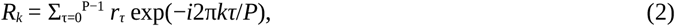

is evaluated at the first harmonic (k = 1, the stimulus frequency itself, the signature of a unit that brightens once and dims once per cycle) and second harmonic (k = 2, twice the stimulus frequency, the signature of a biphasic, frequency-doubling unit). Normalizing c_1_ = R_1_/P and c_2_ = R_2_/P gives amplitudes and phases:

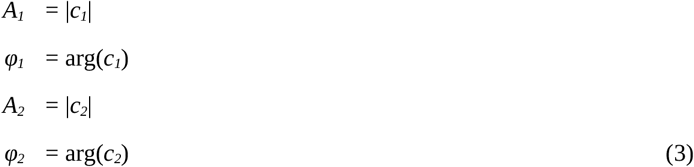

Units with ρ = A_2_/A_1_ ≥ θρ (θρ = 0.6) are routed to the biphasic branch (k = 2); otherwise they are treated as ON/OFF candidates (k = 1); the same numerical-stability constant ε is added to A_1_ in this ratio, so a vanishingly small A_1_ routes the unit to the biphasic branch by construction rather than causing a division error. Before a label is accepted, each candidate must pass three reliability gates evaluated at the relevant harmonic:

i. a signal-to-noise ratio

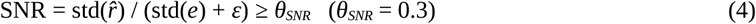

where *r6* is the response reconstructed from only the relevant harmonic (and its conjugate), *e* = *r* − *r6* is the residual, and *ε* = 1×10^−9^ is a fixed numerical-stability constant added to this and all other such denominators throughout the pipeline to avoid division by zero;
ii. a per-cycle amplitude floor,

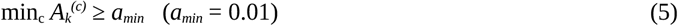

requiring the harmonic to be present in every analyzed cycle, not just on average; and
iii. a phase-jitter limit,

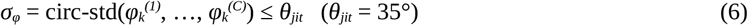

requiring the response to occur at a consistent phase of the stimulus from cycle to cycle. Units failing any gate are marked Unclassified, with the failing criterion recorded.

ON/OFF-branch units are assigned a polarity from the peak timing of the first-harmonic reconstruction

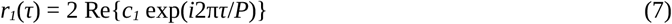

where the OFF window spans the central half of the cycle (*τ* ∈ [0.25*P*, 0.75*P*), corresponding to 125-375 ms of the 500 ms cycle at the 10 ms bin width and *P* = 50 bins/cycle used here) and the ON window is its complement, wrapping across the cycle boundary (*τ* ∈ [0.75*P, P*) ∪ [0, 0.25*P*), 375-125 ms). With P = 50, the implemented OFF window is bins 12-37 (120-380 ms) and the ON window is bins 38-49 and 0-11. Cycles whose reconstructed peak falls in the OFF window vote OFF and cycles whose peak falls in the ON window vote ON, and the unit is labeled by majority vote across the five analyzed cycles. Because the number of cycles is odd, a tie cannot occur; the code retains a tie-break by the sign of the mean first-harmonic real-part (in-phase) projection (ON if ≥ 0). A kinetic label (sustained/transient) is assigned from the effective width 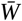, the mean over cycles 2-6 of a per-cycle width computed from the baseline-adjusted response in the unit’s polarity window,

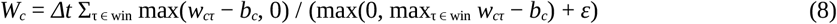

Here, *w*_*cτ*_ is the response of cycle *c* at within-cycle bin *τ* after mean-centering the cycle and orienting its dominant deflection to be positive within the assigned polarity window, “win” (OFF, bins 12-37; ON, bins 37-49 and 0-12, so the two windows share their edge bins); *b*_*c*_ is the median response of that cycle outside the window; and *Δt* is the bin width (10 ms for the recordings analyzed here, see Preprocessing above). 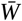 is the mean of *w*_*c*_ over cycles 2-6. Each per-cycle width is the duration of a rectangle with the same area and peak as that cycle’s baseline-adjusted response (ε = 1×10^−9^ as above). 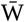 is compared against a threshold Wthr: sustained if 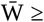 Wthr, transient if 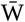 < Wthr. A unit that passes the polarity gates but whose per-cycle peak falls below peak_floor in any analyzed cycle is dropped from the kinetic (transient/sustained) labeling step entirely; it retains its ON/OFF polarity label but receives no kinetic subtype. Width is computed only for units whose peak response within the expected window reaches at least a minimum z-score, peak_floor = 0.5, so that an effective width is not estimated from a peak too small to be reliably localized. Default parameter values are summarized in Table 1.

### Hidden Markov model for population activity

The binary population activity is modeled as generated by a sequence of latent states *Z*_*1:T*_ = {*z*_*1*_,…,*z*_*T*_}, *z*_*t*_ ∈ {1,…,*K*}, with the first-order Markov property and the standard HMM emission-independence assumption. The model is specified by an initial state distribution *π*_*k*_ = *P*(*z*_*1*_=*k*), a transition matrix *A*_*ij*_ = *P*(*z*_*t*_=*j* | *z*_*t−1*_=*i*), and an emission distribution *B*_*j*_(*x*_*t*_) = *P*(*x*_*t*_ | *z*_*t*_=*j*). The joint probability of hidden states and observations is

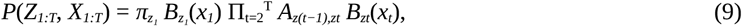

and the most likely hidden-state sequence given the observations,

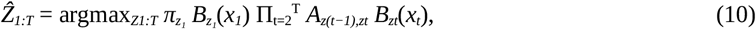

is obtained with the Viterbi algorithm after parameter estimation.

Parameters *π, A*, and the emission parameters are estimated by expectation-maximization (Baum et al., 1970; Rabiner, 1989). In the E-step, the forward-backward recursion computes

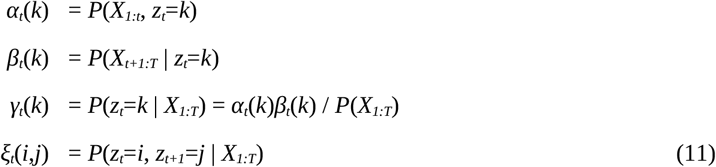

(state occupancy posterior and pairwise posterior, respectively). In the M-step,

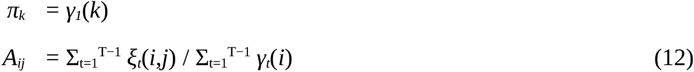

and the emission parameters are updated according to the Chow-Liu tree emission model below. Because the HMM log-likelihood is not concave, each model is fit from 5 random initializations (seeds 0-4) for up to 60 iterations or until the absolute change in training log-likelihood per bin between successive EM iterations falls below Δ = 5×10^−4^, and the solution with the highest training log-likelihood is retained. (During the cross-validated hyperparameter search, a 30-iteration cap and a tolerance of Δ = 1×10^−4^ are used; the values above apply to the final refit. The stopping rule does not require the likelihood to increase monotonically.) Because the emission update is covariance-thresholded, it is a regularized update rather than an exact maximum-likelihood M-step.

### Chow-Liu tree emission model

Directly modeling *P*(*x*_*t*_ | *z*_*t*_=*k*) over *N* binary units requires 2^*N*^ − 1 parameters per state, which is infeasible to estimate. PHASOR instead approximates each state-conditional emission distribution by a Chow-Liu tree (Chow and Liu, 1968), which captures the strongest pairwise dependencies among units, which account for much of the structure in real neural population activity (Schneidman et al., 2006), with only *N* − 1 additional parameters per state. For state *k*, weighted pairwise counts are accumulated using the posterior responsibilities *γ*_*t*_(*k*),

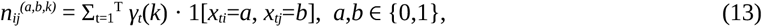

giving pseudo-count-smoothed pairwise and marginal probabilities 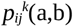 and 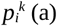 (symmetric pseudocount α = 0.5, added to each of the two marginal outcomes and to each of the four joint outcomes, so that every smoothed table is strictly positive; α is held fixed rather than searched). The within-state pairwise covariance is then regularized by an absolute soft-threshold of size η ≥ 0, replacing each covariance by sign(c) max(|c| − η, 0), so that any pair whose weighted covariance under that state does not exceed η is treated as exactly independent. Each pairwise table is rebuilt from the common node marginals together with the thresholded covariance, clipped so that all four cell probabilities remain strictly positive. Because every edge table is by construction consistent with the same node marginals, the Chow-Liu factorization below is an exactly normalized probability distribution and requires no additional partition term. The mutual information between units i and j under state k, computed from these regularized tables,

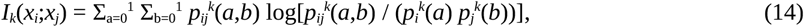

is used as an edge weight to build a maximum-spanning tree *T*_*k*_ over the *N* units, and the emission distribution is approximated by the Chow-Liu factorization

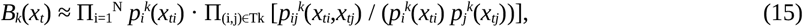

with log-emission score

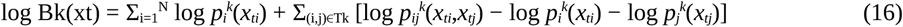

where the second sum runs over the N − 1 edges of that state’s maximum-spanning tree. Because each edge table shares the node marginals used in the first sum, this score is an exactly normalized log-probability at every η ≥ 0, so held-out log-likelihoods are directly comparable across the candidate (K, η) grid. Setting η = 0 leaves the α-smoothed covariance unthresholded, while increasing η progressively prunes weakly co-varying edges toward the independent-Bernoulli limit. η is selected per recording by cross-validation from the grid η ∈ [0.0005, 0.002, 0.005], a grid centered on small values under the assumption that pairwise covariances between simultaneously recorded units are comparatively weak; this assumption is addressed further as a limitation in the Discussion. All ten retinal recordings selected η = 0.005, the largest value in the grid, with α fixed at 0.5 (Table 2).

### Model selection

The number of states K and the covariance threshold η are selected by forward-chaining cross-validation over 3 folds, with the pseudocount held fixed at α = 0.5: the training data (repetitions 1-10, concatenated in presentation order and fit as one sequence without resets; 30,000 bins for WT1 run 1) are split so that the model is fit on the first 40%, 60%, or 80% of bins and scored by log-likelihood per bin on the following 20%. Repetitions 11-15 are excluded from fitting and model selection and are decoded separately. This is repeated across a set of candidate K chosen per dataset to bracket the expected elbow (K = [5, 6, 7, 8, 9] for the K = 7 synthetic datasets, K = [8, 9, 10, 11, 12] for the K = 10 synthetic dataset, and K = [3, 6, 9, 12, 15] for the retinal recordings reported here), over η ∈ [0.0005, 0.002, 0.005], and over [0, 1, 2, 3, 4] random seeds. For each candidate K the η value with the highest mean validation log-likelihood per bin is retained, ties being broken by the lower standard deviation across folds and seeds. Validation log-likelihood typically increases with K without a clear maximum, favoring progressively more fragmented, over-segmented models (Li and La Camera, 2025; Pohle et al., 2017); PHASOR therefore selects K using the following elbow procedure, applied to the marginal gain ΔyK = yK+1 − yK in the best validation score yK at each candidate K (Algorithm 1):

#### Algorithm 1.

Elbow rule for model-order selection

1. If there are fewer than three candidates, or all gains are equal, select the K with the largest validation score. Otherwise, if the final marginal gain is at or below the 20th percentile of all observed gains, select the second-to-last candidate *K*.
2. Otherwise, define a plateau scale as 1.5 × the median of the last two marginal gains, and select the smallest *K* beyond which all remaining gains stay at or below this scale.
3. If neither rule identifies a plateau, select the K with the largest validation score. In the event of an exact tie in validation score, the smallest tied K is selected (a consequence of how the fallback rule scans candidates in ascending K order).

Under the selected hyperparameters, the model is re-fit from 5 random initializations (seeds 0-4) and the fit with the highest training log-likelihood is retained.

### State summary and comparison across recordings

Once fit, PHASOR reports each state’s occupancy, per-unit firing probability, entropy relative to a combinatorial upper bound, self-transition strength, and off-diagonal transition entropy, together with the Viterbi-decoded state sequence for training and held-out repetitions. Mean dwell time for state k is the mean length of consecutive runs of k in the Viterbi-decoded sequence multiplied by the 10 ms bin width (not the model-implied value derived from the self-transition probability); off-diagonal transition probabilities are renormalized (self-transitions removed, rows re-summed to 1) before computing off-diagonal entropy. A unit is called active within a state if it fired at least one spike in any bin of the analyzed window. An optional module, not used for any result reported here, compares fitted states across recordings or preparations using posterior-weighted similarity. Each state is represented by three feature blocks. The emission block contains the 25th, 50th, and 75th percentiles of the *ϕ* correlations across its Chow-Liu edges, where

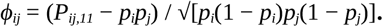

The temporal block contains the self-transition probability *A*_*ii*_, full-row transition entropy *H*_full_, calculated from the complete transition-probability row, and off-diagonal transition entropy *H*_off_, calculated after removing the self-transition and renormalizing the remaining probabilities. Each temporal feature is centered by its pooled median and divided by its pooled interquartile range across the calibration recordings. The stimulus-overlap block contains ten binary codes based on a relative-occupancy threshold of 0.60, with present and absent epochs coded as 0 and 1, respectively. The emission, temporal, and overlap blocks are divided by 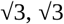, and 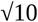, respectively. Euclidean distances are calculated separately for each block between every cross-recording state pair and converted to blockwise similarities using Gaussian kernels with robust, globally calibrated distance scales; the three blockwise similarities are then averaged to obtain state-pair similarity. Posterior state probabilities are averaged across all 15 stimulus repetitions. At each stimulus-aligned time bin, every state-pair similarity is weighted by the product of the corresponding posterior probabilities, and the resulting values are summed across state pairs and averaged over time to obtain the recording-level similarity. This procedure does not require discrete state matching, a distance threshold, or identical numbers of states across recordings.

### Simulated validation data

To validate the HMM fitting and model-selection procedures against known ground truth, synthetic population activity was generated using a chain-structured (tree) conditional Bernoulli cascade with hop-distance-decaying pairwise dependencies of the kind a Chow-Liu fit is designed to recover (see Extended Data Fig. 2-1), using the same 166-event, 828.5 s stimulus protocol as the retinal recordings (Extended Data Fig. 2-1A), for a fixed generative state count (K = 7: one baseline (contrast 0%) state and six stimulus-locked, phase-tuned states) and, separately, K = 10 (one baseline (contrast 0%) plus nine active states); 240 simulated units (80 ON-tuned, 80 OFF-tuned, 80 biphasic-tuned, fixed by unit index; assignment of units to assemblies and the order of each chain are drawn at random for each seed). A second, fully deterministic generator, in which assembly units fire exactly once per bin within their assigned block and non-selective units follow an independent Poisson process, served only as a basic pipeline sanity check and is not part of the validation results reported here; all reported validation results use the stochastic, phase-tuned tree generator described above. Ten independent random seeds (0, 1, 2, 3, 4, 5, 6, 7, 8, and 9500) were simulated at K = 7 and passed through the full PHASOR pipeline, including forward-chaining cross-validation over a candidate range of K = [5, 6, 7, 8, 9] and the elbow rule for K selection; a ground-truth alignment between generative and decoded states was available for all 10 seeds. The K = 10 dataset (seed 9501; Fig. 2D-F) was fitted separately, over a candidate range of K = [8, 9, 10, 11, 12] (see Results for a comparison of the two).

**Figure 2.**
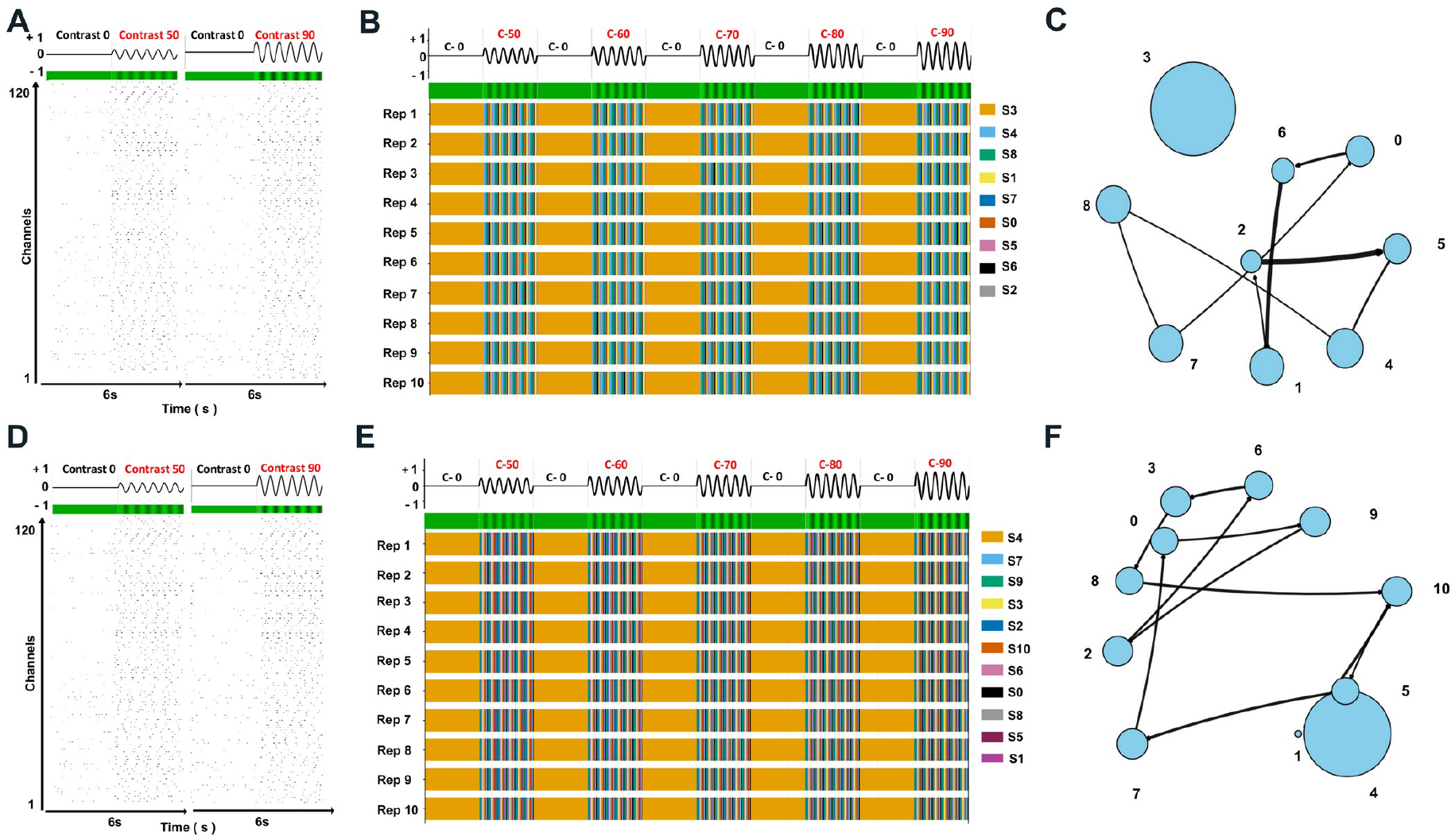
Application of PHASOR to synthetic datasets generated at two distinct state counts. (A-C) Dataset generated using the synthetic data pipeline (Extended Data Fig. 2-1) with K-generative = 7 (seed 9500; one baseline state plus six stimulus-locked active assembly states, per Materials and Methods). (A) Raw population raster (120 channels) across the C-50-C-90 contrast protocol. (B) Globally optimal state sequence decoded via the Viterbi algorithm for repetitions 1-10; PHASOR selected a 9-state model (S0-S8) for this dataset. (C) Transition-structure graph of the PHASOR-fitted 9-state model (generative K = 7); node size is proportional to state occupancy, edge width is proportional to transition probability, edges are directional (drawn separately for each ordered state pair i -> j), and only edges with P(i,j) exceeding a threshold of 0.05 are displayed. (D-F) Dataset generated using the same synthetic data pipeline (Extended Data Fig. 2-1) with K-generative = 10 (seed 9501; one baseline state plus nine stimulus-locked active assembly states, following the same convention as the K = 7 case). (D) Raw population raster (120 channels) across the C-50-C-90 contrast protocol. (E) Globally optimal state sequence decoded via the Viterbi algorithm for repetitions 1-10; PHASOR selected an 11-state model (S0-S10) for this dataset. (F) Transition-structure graph of the PHASOR-fitted 11-state model (generative K = 10), with node size proportional to state occupancy and edge width proportional to transition probability. The rasters in (A) and (D) are shown by the 120 recording channels, not by the 240 individual synthetic units.

### Statistical analysis

Recovery of the known generative state assignment in the K = 7 synthetic validation data (see Simulated validation data above) was quantified using the adjusted Rand index (ARI; Hubert and Arabie, 1985), a chance-corrected, permutation-invariant measure of agreement between two categorical labelings that does not require the generative and decoded state indices to be aligned. For each of the ten seeds, ARI was computed (scikit-learn adjusted_rand_score) between the per-bin generative phase-state label and the per-bin Viterbi-decoded state label, separately for the training repetitions (1-10) and the held-out repetitions (11-15), and within each of those for all active-stimulus bins (contrast > 0), for the 50%-contrast block alone, and for the 90%-contrast block alone; baseline (contrast = 0) bins were excluded from these active-period comparisons because the generative process defines a single, trivial state throughout baseline, for which ARI is not informative by construction. Results are reported as the range and mean across the ten seeds (Statistical Table); ranges are observed minima and maxima, not confidence intervals, and ARI is an agreement measure rather than a hypothesis test. For the confusion matrix (Fig. 3D), decoded states were matched one-to-one to generative states with the Hungarian algorithm (Kuhn, 1955; Munkres, 1957); a decoded state without a generative partner remained unmatched.

**Figure 3.**
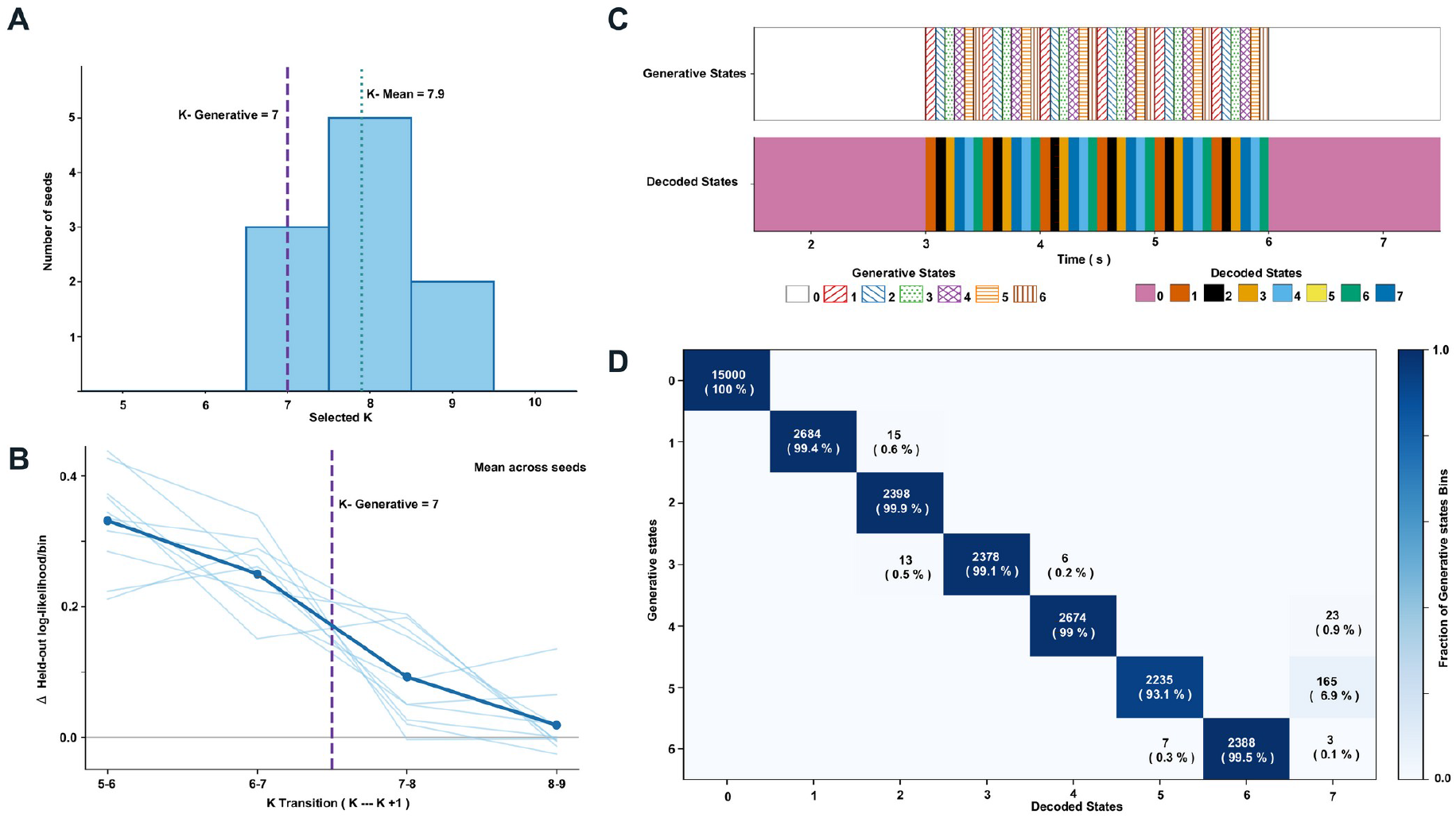
Model-order selection and recovery of generative states in synthetic data with known ground truth. (A) Distribution of PHASOR’s selected model order (K*) across n = 10 independently seeded synthetic datasets generated with a true generative order of K-generative = 7 (dashed line); mean selected order across seeds, K-mean = 7.9 (dotted line). (B) Marginal gain in cross-validated log-likelihood per bin (ΔLL) for each successive candidate-order transition (K → K+1); thin light-blue lines show individual seeds, the bold dark-blue line shows the across-seed mean, and the dashed line marks the true generative order transition. (C) Representative single-seed (seed 0, for which K* = 8) comparison of the ground-truth generative state sequence (top; white = baseline/state 0, hatched colors = states 1-6) and PHASOR’s Viterbi-decoded sequence (bottom; solid colors S0-S7) over a representative window. (D) Confusion matrix for the same representative seed, quantifying the correspondence between generative states (rows, 0-6) and decoded states (columns, 0-7); values are training-bin counts (including baseline) with the row-normalized percentage in parentheses, and the color scale indicates the fraction of a generative state’s bins assigned to each decoded state.

### Code and data availability

The PHASOR source code is available at https://github.com/Phasor-population-hmm/PHASOR. PHASOR’s analysis backend is implemented in Python (≥3.10) and its browser-based interface in Next.js (Node.js ≥20); the code is released under the MIT License. The repository includes the complete source for both interfaces, a README with installation instructions, the per-stage command-line invocations described in Software overview above, a dependency specification (requirements.txt), and example data for the ten recordings in Table 2. The analyses reported here were run on Ubuntu 20.04.6 LTS (Intel Xeon W-2245) and macOS Tahoe 26.0 (Apple M3 Pro).

## Results

### Pipeline overview

Fig. 1 traces PHASOR from raw recording to state-resolved interpretation: spikes are aligned to the stimulus clock, each unit receives a Fourier response class, a Chow-Liu-tree HMM is fit with its state count chosen by cross-validation, and decoded states are cross-referenced with those classes. It is applied below to synthetic data, then to the ten recordings in Table 2.

### Validation on simulated data with known ground truth

Applying the pipeline to synthetic datasets built at two generative state counts gave a consistent pattern (Fig. 2). At K-generative = 7, the raw population raster across the C-50 to C-90 protocol (Fig. 2A) was decoded into a 9-state model (S0-S8) recurring consistently across repetitions (Fig. 2B), with the fitted transition structure concentrating occupancy and transition probability on a small subset of states (Fig. 2C). At K-generative = 10, the same settings returned an 11-state model (S0-S10) with a denser transition graph (Fig. 2D-F).

The same generative process (K = 7, 240 units) was then resimulated across ten seeds. The elbow rule, applied to forward-chaining cross-validated log-likelihood, selected K* between 7 and 9 (mean 7.9), matching the generative K exactly in three of ten seeds and never falling below it (Fig. 3A); the mean marginal gain in cross-validated log-likelihood per bin fell from 0.25 at the 6 to 7 transition to 0.09 at 7 to 8 and 0.02 at 8 to 9, the plateauing rather than peaking that motivates the elbow criterion (Fig. 3B). For seed 0 (K* = 8), decoded and generative states corresponded nearly one to one: 99.2% of training bins fell on the Hungarian-matched diagonal and the single unmatched decoded state took 0.7% of bins (Fig. 3C,D), and agreement was equally high on held-out repetitions (ARI 0.988 over active-stimulus bins, against 0.978 on training). Across the ten seeds, ARI over active-stimulus bins was 0.868-0.994 (mean 0.963) on training and 0.867-0.992 (mean 0.965) on held-out repetitions ^a^, and was similar within the 50% and 90% contrast blocks^b,c^ (Statistical Table).

### Demonstration on ex vivo retinal recordings

In the WT1 run 1 recording, mean firing rate was flat between 0 and 50% contrast and rose steeply above it (Fig. 4A,B). Splitting units by Fourier polarity separated cycle-averaged responses into ON, OFF, and biphasic groups with distinct phase relationships to the stimulus (Fig. 4C) and distinct contrast-response curves, ON units reaching the highest rates at 90% contrast (Fig. 4D); the kinetic label further divided ON and OFF into transient and sustained subtypes differing in response width and in position relative to the ON and OFF stimulus windows (Fig. 4E). Active-unit counts across twelve stimulus-cycle phase partitions increased between 50% and 90% contrast while subtype phase ordering was preserved (Fig. 4F).

**Figure 4.**
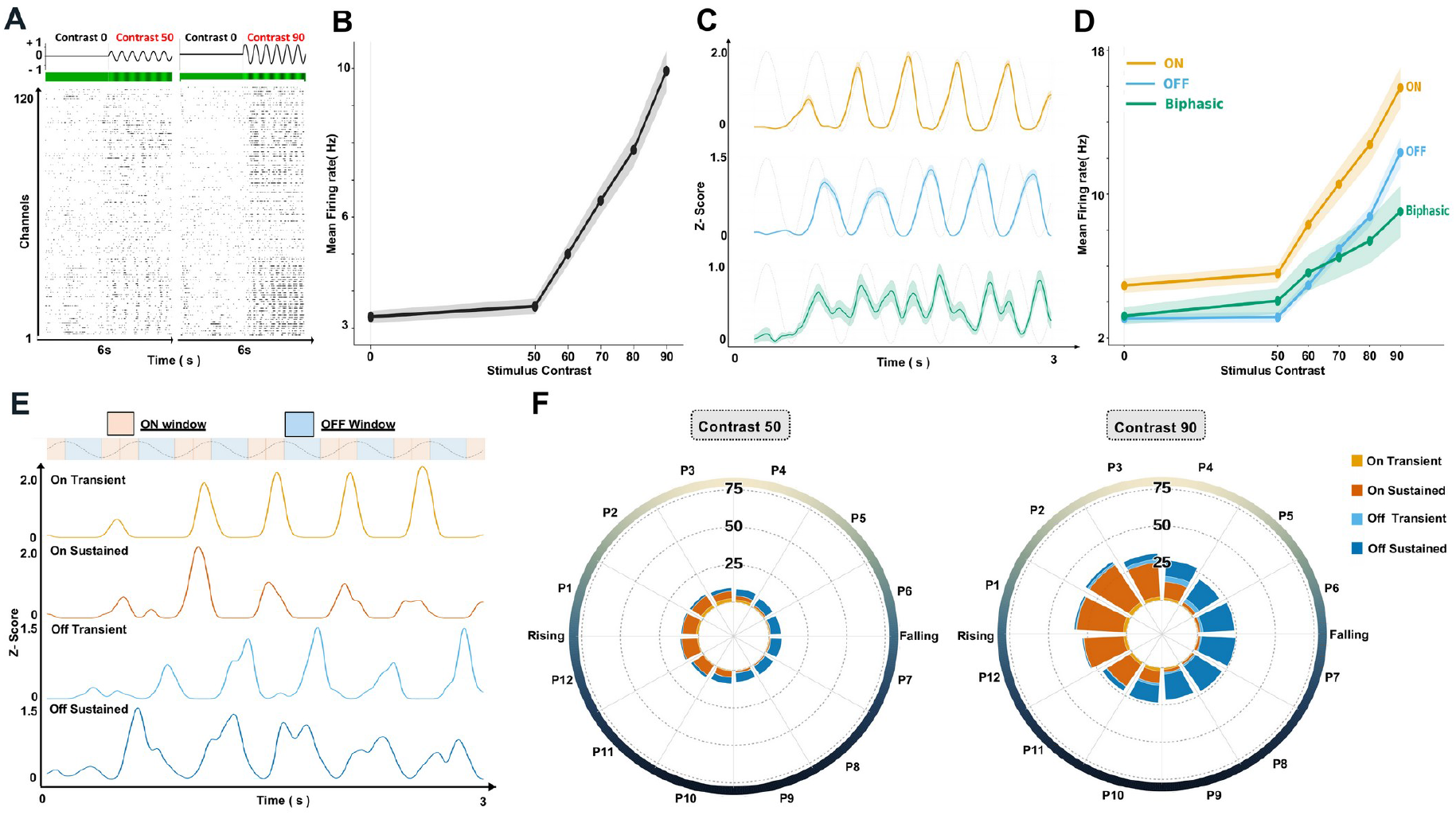
Contrast- and response-class-dependent response dynamics in WT1 run 1. (A) Representative population raster (120 channels) during the 0-50% and 0-90% contrast transitions. (B) Mean firing rate (± SEM) as a function of stimulus contrast, pooled across all 276 units in this recording. (C) Mean z-scored response (± SEM) for ON, OFF, and biphasic cells aligned to stimulus phase. (D) Mean firing rate (± SEM) versus contrast for the ON, OFF, and biphasic subpopulations. (E) Z-scored response traces for the four On/Off subtypes (On-transient, On-sustained, Off-transient, Off-sustained); shaded background bands mark the “ON window” and “OFF window” used to assign phase labels. (F) Polar histograms of active-cell counts by response subtype across 12 stimulus-phase partitions (P1-P12, spanning the rising and falling half-cycles) at 50% (left) and 90% (right) contrast. These panels show one retina; SEM summarizes within-recording variability. ON/OFF/biphasic and transient/sustained are functional labels assigned by the classifier described in Materials and Methods.

On real data the marginal validation gain again decreased monotonically with candidate model order without a maximum (Fig. 5A, top; cf. Fig. 3B), and the elbow rule selected a 12-state model (276 units; η = 0.005, α = 0.5; Table 2) whose decoded sequences recurred across repetitions and realigned at each contrast block (Fig. 5A, bottom). Decoded states differed in occupancy and dwell time: State 2 held roughly a quarter of all bins with the longest mean dwell time (∼340 ms), State 5 was rarely visited (∼1%), and the remaining states held ∼5-11% of bins with dwell times of ∼65-190 ms (Fig. 5B). Transition-matrix entropy stayed below the theoretical maximum (∼3.6 bits) for every state, and off-diagonal entropy (up to ∼2.3 bits) exceeded full-row entropy for most states, indicating self-transition-dominated rather than unstructured switching (Fig. 5C). Cross-referencing states with per-unit labels resolves transitions by subtype: at one transition the preceding state was OFF-dominated (29 of 64 active units, predominantly Off-sustained) and the following state ON-dominated (21 of 51, predominantly On-sustained), so the phase change coincided with a subtype shift (Fig. 5D).

**Figure 5.**
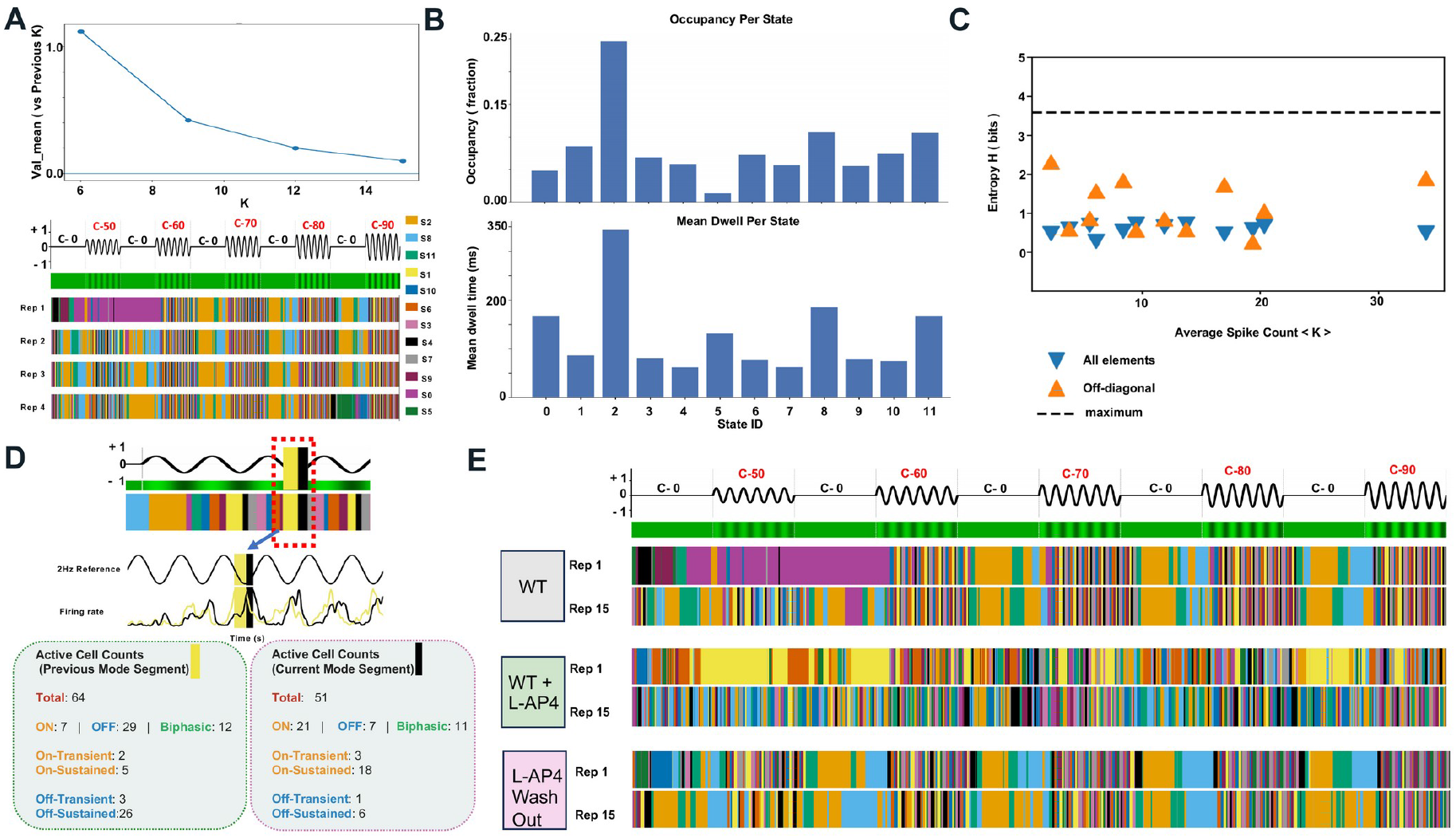
Model-order selection diagnostics and repeated analysis across pharmacological conditions. (A) Top, mean validation criterion (cross-validated likelihood gain relative to the previous K) versus candidate model order K for a representative recording (WT1, run 1). Bottom, corresponding Viterbi-decoded state sequences (repetitions 1-4) for the same recording across the stimulus protocol, colored by decoded state. (B) Occupancy (fraction of total bins, top) and mean dwell time (mean length of decoded runs, ms; bottom) for each decoded state (State ID 0-11) of the model. (C) Transition-matrix entropy (bits) versus average spike count (⟨K⟩) for each state of the model, shown separately considering all transition-matrix elements for that state’s row (blue, downward triangles) or only its off-diagonal, state-switching elements (orange, upward triangles). The dashed line marks log2(12), the maximum for the full transition row. Off-diagonal probabilities are renormalized over 11 possible destinations, whose maximum entropy is log2(11). (D) Example of a decoded transition between two consecutive states occurring at different phases of the stimulus cycle in the representative recording: stimulus/firing-rate trace (top) with the analyzed transition window highlighted, and active-cell counts by ON/OFF/biphasic class and by On/Off-transient/sustained subtype, tallied separately for the state segment immediately preceding and following the transition. The preceding state is dominated by OFF cells (29 of 64 active units, predominantly Off-sustained), while the following state is dominated by ON cells (21 of 51 active units, predominantly On-sustained), showing that the shift in phase between the two states coincides with a shift in which cell subtype dominates the active population. Units not shown in these dominant-subtype counts fall into the other ON/OFF/biphasic categories or are Unclassified (see Materials and Methods for the classifier’s reliability gates). (E) Viterbi-decoded state sequences (repetitions 1 and 15) across the whole stimulus protocol for three sequential pharmacological conditions applied to the same retina preparation: wild-type (WT), WT + L-AP4 (an mGluR6 agonist (Slaughter and Miller, 1981; Nakajima et al., 1993) that suppresses ON bipolar cell signaling), and after L-AP4 washout. Each condition was fit as an independent HMM, so identical state colors across panels do not by themselves imply the same underlying state. Related outputs are shown in Extended Data Figures 5-1 (ON-labeled peak latency under L-AP4), 5-2 (no-projection recording WT3), and 5-3 (rd1 recording rd1-1).

Applying the pipeline across a pharmacological manipulation of the same preparation demonstrates the decoded representation’s sensitivity to a known perturbation (Fig. 5E). Under L-AP4, which suppresses ON-bipolar cell signaling, the decoded state sequence differed visibly from the pre-treatment condition, and the washout sequence appeared closer to it (not quantified); the three conditions were fit as independent HMMs, so identical state colors do not by themselves indicate correspondence. A second wild-type retina (WT2) was taken through the identical protocol with the same settings (Table 2). A peak-latency analysis of ON-cell responses in the same preparation showed a small delay of the mean ON-cell peak under L-AP4 at both 50% (+38.0 ms, +27.4°) and 90% (+26.0 ms, +18.7°) contrast (Extended Data Fig. 5-1); because the classifier is re-run per condition and the compared groups differ in size, this is an independently averaged population comparison, not a within-cell one.

Two controls bound these interpretations. With the same protocol run in the dark and no stimulus projected, mean firing rate stayed at roughly 2-4 Hz across every nominal contrast label, against a rise to ∼10 Hz at 90% contrast when the stimulus was delivered; decoded states were scattered in time, with several repetitions held in a single state and no realignment at contrast-block boundaries (Extended Data Fig. 5-2A-C). Full-row transition entropy was lower than in the wild-type fit (∼0.2-0.4 bits against ∼0.3-0.9 bits), and no off-diagonal transition exceeded the display threshold (Extended Data Fig. 5-2D,E). The complementary comparison keeps the stimulus but uses a photoreceptor-degenerate retina: *rd1*-1 recorded under the identical projected protocol, also showed no contrast recruitment, in a different experimental context. Its mean firing rate was already near the level the wild-type recording reached only at 90% contrast and varied little across the range (Extended Data Fig. 5-3A,B); its fitted 12-state model used all of its states, yet the decoded sequences did not realign at contrast blocks (Extended Data Fig. 5-3C). Its off-diagonal transition entropy (∼1.5-3 bits) was high for every state, whereas the wild-type fit included states with more selective exits (Extended Data Fig. 5-3D,E; Fig. 5C).

### Statistical Table

Recovery of the K = 7 synthetic ground-truth state assignment by the fitted HMM/Viterbi decoder (Simulated validation data; Statistical analysis, Materials and Methods).

| Data structure | Type of test | Result (observed range and mean across seeds; not a confidence interval) |
|---|---|---|
| <sup>a</sup> Generative vs. Viterbi-decoded state label, per bin; all active-stimulus bins (contrast > 0); n = 10 independent K = 7 synthetic seeds | Adjusted Rand Index<br>(Hubert and Arabie, 1985) | Training repetitions, range 0.868 to 0.994, mean 0.963; held-out repetitions, range 0.867 to 0.992, mean 0.965 |
| <sup>b</sup> Generative vs. Viterbi-decoded state label, per bin; 50%-contrast block only; n = 10 seeds | Adjusted Rand Index | Training repetitions, range 0.870 to 0.991, mean 0.961; held-out repetitions, range 0.868 to 0.989, mean 0.961 |
| <sup>c</sup> Generative vs. Viterbi-decoded state label, per bin; 90%-contrast block only; n = 10 seeds | Adjusted Rand Index | Training repetitions, range 0.872 to 0.996, mean 0.964; held-out repetitions, range 0.872 to 0.998, mean 0.969 |

## Discussion

On synthetic data with a known generative state count, the elbow-selected K matched or modestly overshot the generative K (exact recovery in three of ten seeds; mean K* = 7.9), while the decoded states recovered the generative state sequence with high fidelity across all ten seeds (Results; Statistical Table). That dissociation is the central methodological point of the validation: a model-order criterion can miss the generative state count by one or two states and still return a state sequence that is almost perfectly aligned with the generative one, so sequence recovery and exact recovery of K provide complementary validation criteria. Recovery was similar at the lowest and highest contrast tested (mean ARI 0.961 at 50% and 0.964 at 90% contrast on the training repetitions) and similar again on the held-out repetitions, indicating that the decoded states tracked the generative sequence across the contrast range of the generative model.

Applied to retinal recordings, the same kinds of outputs are produced. In the representative wild-type recording the selected model used twelve states whose occupancies spanned more than an order of magnitude, whose dwell times ranged from tens to hundreds of milliseconds, and whose transition structure was dominated by self-transitions rather than by unstructured switching. Because every unit also carries an independent Fourier response label, each state can additionally be read in terms of the cells active within it, and the two descriptions are informative together in a way neither is alone: a transition that appears in the state sequence as a change in stimulus phase appears in the unit labels as a handover from an Off-sustained-dominated to an On-sustained-dominated active population. The pharmacological series makes the same point on a manipulation whose target is known. Suppressing ON bipolar cell transmission with L-AP4 changed the decoded sequence visibly, with the washout sequence appearing closer to the pre-treatment one (not quantified), and an independent peak-latency measurement on the same preparation showed the ON-cell response delayed by tens of milliseconds.

The two control recordings are included for a specific reason. A state description of this kind can be produced from any spike train, and neither its existence nor its apparent structure demonstrates that it reflects the stimulus. Withholding the projection while keeping the protocol timing removes both the contrast-dependent rate increase and the realignment of states at block boundaries, indicating that the protocol timing alone does not produce them (Extended Data Fig. 5-2). Recording an *rd1* retina, which is photoreceptor-degenerate and also differs in strain and animal, under the identical projected stimulus produces a different failure mode: activity is high and essentially contrast-invariant from the outset, and the decoded states, although numerous, do not lock to the stimulus (Extended Data Fig. 5-3). The two controls separate two confounds that either one alone would leave entangled, and together they define what the stimulus-locked structure in the wild-type recordings is being contrasted against. Both are single recordings and are presented as qualitative controls rather than as a powered genotype comparison.

PHASOR’s modular architecture allows its emission model, classification thresholds, and model-selection criteria to be adjusted or extended as additional data and use cases arise. Because every stage writes its output in a fixed, documented format, individual stages can be re-fit or replaced without repeating earlier steps, and the same procedure can be applied consistently across recordings, as it was across the ten recordings reported here (Table 2).

The main limitations of this work follow from the assumptions built into the model. The emission model assumes that dependencies between simultaneously recorded units are weak and adequately captured by a small number of pairwise terms, and the covariance threshold was restricted to small values for that reason; populations carrying materially stronger dependencies may not be well served by that choice. The hidden Markov model further assumes that the population state in each time bin depends only on the state in the bin before it, so slower history effects and longer-range dependencies are not represented. The synthetic data used for validation were generated with a tree-structured, hop-distance-decaying dependency so that the simulated population would carry correlations of the kind the emission model is meant to describe; a generator without such correlations would not exercise that part of the model, and the present validation therefore does not probe performance when the true dependency structure is very different. The synthetic states are prescribed phase blocks with contrast- and gain-dependent emissions, so the generative K is not a minimal stationary-HMM order; the validation tests label recovery. The model-order selection rule is a heuristic, not a guarantee: in the synthetic data it never fell below the generative count but often exceeded it, and its choice depends on the candidate grid. With the retinal grid (3, 6, 9, 12, 15), a steadily shrinking gain returns the second-to-last candidate (K = 12, selected in nine of ten recordings; WT2 run 2 selected K = 15), and every retinal fit selected the largest tested η (Table 2), so neither K nor η should be read as an optimum. The Fourier classifier describes each unit through the first two harmonics of a single periodic stimulus, so units whose responses are weak or poorly captured by those harmonics are left Unclassified, and the resulting labels describe responses to this stimulus rather than cell identity in general. Finally, the retinal material is a demonstration of the workflow rather than a biological study: the main-text figures come from one representative recording, the pharmacological series includes two retinas, whereas each control is represented by one retina (Table 2), and the dispersion shown in the figures is within-recording variability rather than biological replication. Quantitative claims about how these state descriptions differ across animals, genotypes, or drug conditions would require a powered design and are not made here.

## Acknowledgments

ChatGPT (OpenAI) was used to assist with language editing, consistency checks, and submission formatting.

