## Supplemental Figures for "PHASOR: An Open-Source Pipeline for Hidden Markov Model Analysis of Stimulus-Locked Retinal Multielectrode-Array Population Activity"

### PHASOR supplementary figures

#### Figure 2-1

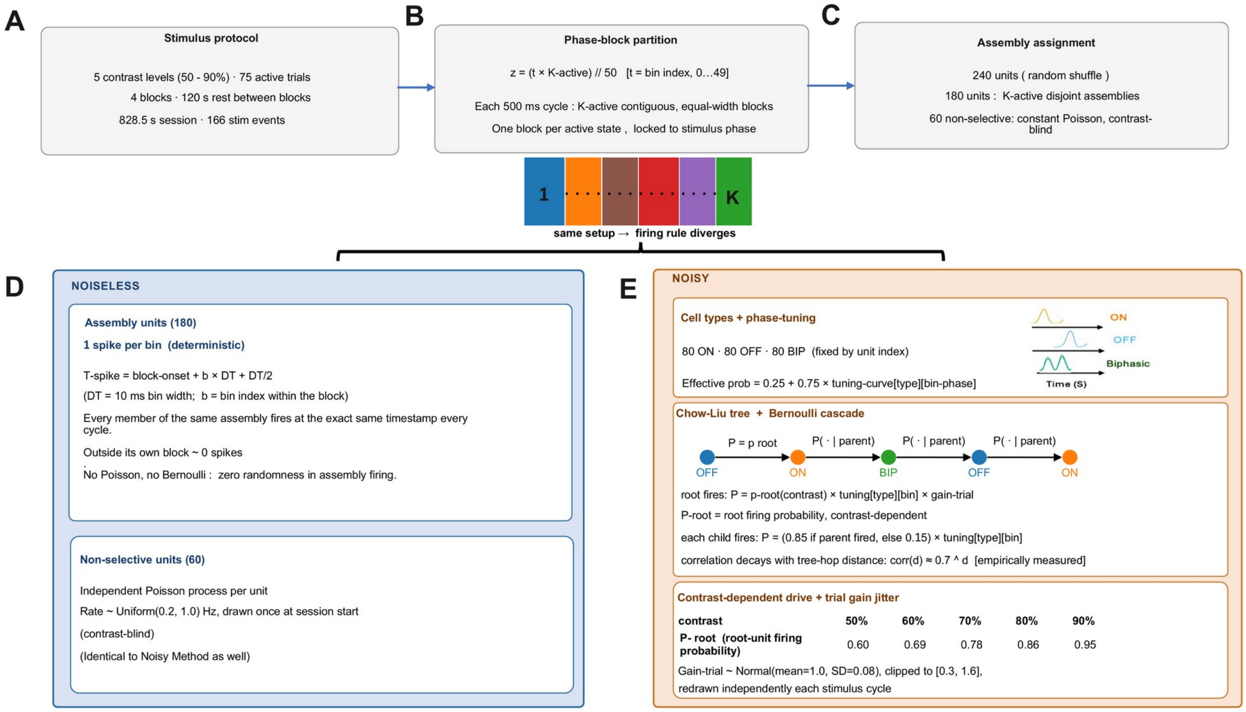


**Figure 2-1.** Synthetic data generation pipeline. Assembly counts in panels C and D illustrate the K = 7 case; the K = 10 allocation is specified below. (A) Stimulus protocol: a 10 s black adaptation period and four 1.04 s integrity flashes, followed by four contrast blocks of 19, 20, 20, and 16 trials (75 trials total; each trial a blank 3 s FFSine at 0% contrast immediately followed by an active 3 s FFSine whose contrast cycles through 50, 60, 70, 80, and 90%), separated by three 120 s rest periods (two consecutive 60 s segments each), then four closing integrity flashes and a final 200 ms black frame; 166 stimulus events, 828.5 s total session duration (active, c >0, time = 225 s, 27.2%; all other time, including blanks, rests, and flashes, counts as baseline, 603.5 s, 72.8%). (B) Phase-block partition: within each 500 ms (50-bin) stimulus cycle, bins are divided into K-active contiguous blocks via z = (t x K-active) // 50 (t = 0-49); at K-true = 7 (K-active = 6) this gives six blocks of 8-9 bins each, identical on every cycle and locked to stimulus phase rather than contrast. (C) Cell-type and assembly assignment: 240 units (80 ON-tuned, 80 OFF-tuned, 80 biphasic-tuned; type fixed by unit index). Within each type, units are randomly permuted for each seed and ⌊60/K-active⌋ are assigned to each assembly, so every assembly mixes all three types: at K-true = 7, 10 per type (30 units) per assembly, leaving 20 per type (60 total) non-selective; at K-true = 10, 6 per type (18 units) per assembly, leaving 26 per type (78 total) non-selective. (D) Noiseless generator (sanity-check only; not used for any reported validation result, see Simulated validation data, Materials and Methods): each assembly unit fires exactly once per bin within its assigned block, at a fixed within-bin offset, with zero trial-to-trial randomness; non-selective units follow the same independent Poisson process as in (E). (E) Noisy generator (tree-structured conditional Bernoulli cascade with phase tuning; used for all reported validation results): within an assembly's active block, assembly members form a chain in random order, and the first (root) unit fires with probability p-root(contrast) x per-cycle gain (product clipped to [0.05, 0.98]) x that unit's own cell-type phase-tuning value at the bin's true phase; p-root increases linearly from 0.60 at 50% contrast to 0.95 at 90% contrast; gain ~ Normal(1.0, SD=0.08), clipped to [0.3, 1.6], redrawn each cycle and shared by all assemblies; each subsequent chain member fires with probability 0.85 (if its parent fired) or 0.15 (if not), again scaled by its own type's tuning value; tuning values are rescaled as 0.25 + 0.75 x tuning curve, so no type's firing probability is ever fully suppressed; because each child depends only on its parent (0.85 vs. 0.15), dependence decays with chain distance d, approximately as 0.7^d for units at full tuning and faster at lower tuning, the hop-distance-decaying structure a Chow-Liu fit is designed to recover. Non-selective units follow an independent Poisson process (rate ~ Uniform(0.2, 1.0) Hz per unit, drawn once at session start, contrast-blind). Abbreviations: p-root, contrast-dependent firing probability of the tree-root unit; gain, per-cycle multiplicative excitability jitter.

#### Figure 5-1

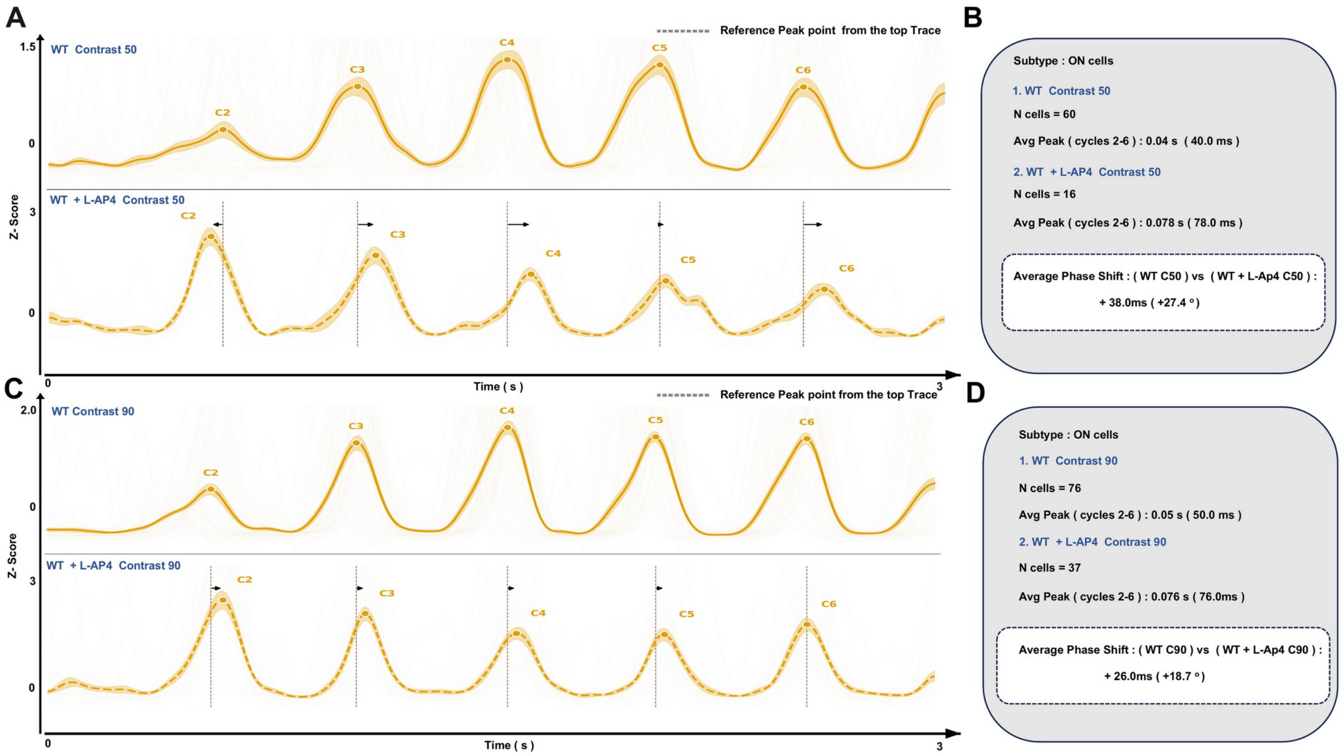


**Figure 5-1.** Peak-to-peak phase analysis of ON-cell responses under ON-pathway blockade in WT1 run 1 and WT1 L-AP4. Mean z-scored ON-cell responses (± SEM) across six stimulus cycles (500 ms each, 2 Hz stimulation) are compared between wild-type (WT) and WT + L-AP4 recordings from the same retina preparation, at 50% (A, B) and 90% (C, D) contrast. (A) Mean ON-cell traces for the 50% contrast condition: WT (top) and WT + L-AP4 (bottom). Filled dots mark the detected peak in each of cycles 2-6 (labeled C2-C6). Cycle 1 is excluded from peak marking and from all quantification, as in the Fourier classifier; only cycles 2-6 are used for analysis. The mean peak latency from the WT trace, averaged across cycles 2-6, is carried down and displayed as a brown dashed reference line at the corresponding time point in every cycle of the WT + L-AP4 trace below; arrows indicate the temporal displacement of each WT + L-AP4 peak relative to this WT reference, with rightward arrows denoting a delayed peak. (B) Quantification for (A): number of ON cells contributing to each trace, mean peak latency within the 500 ms cycle (averaged over cycles 2-6) for WT and WT + L-AP4 separately, and the resulting average phase shift between conditions, reported in milliseconds and as an equivalent phase angle (degrees, scaled to the 500 ms/360° cycle). (C, D) As in (A, B), for the 90% contrast condition: 60 WT and 16 WT+L-AP4 cells contributed to the 50% contrast traces (panel B), and 76 WT and 37 WT+L-AP4 cells contributed to the 90% contrast traces (panel D). The WT and WT+L-AP4 traces are not drawn from the same tracked cohort of cells: because the Fourier classifier is re-run on each condition independently, a given cell can be classified as ON in one condition and Unclassified (or differently labeled) in another, most often due to a weaker signal under L-AP4, so the two traces are independently averaged populations rather than a within-cell comparison. Shading denotes SEM within a single recording and is not a substitute for independent biological (retina- or animal-level) replication.

#### Figure 5-2

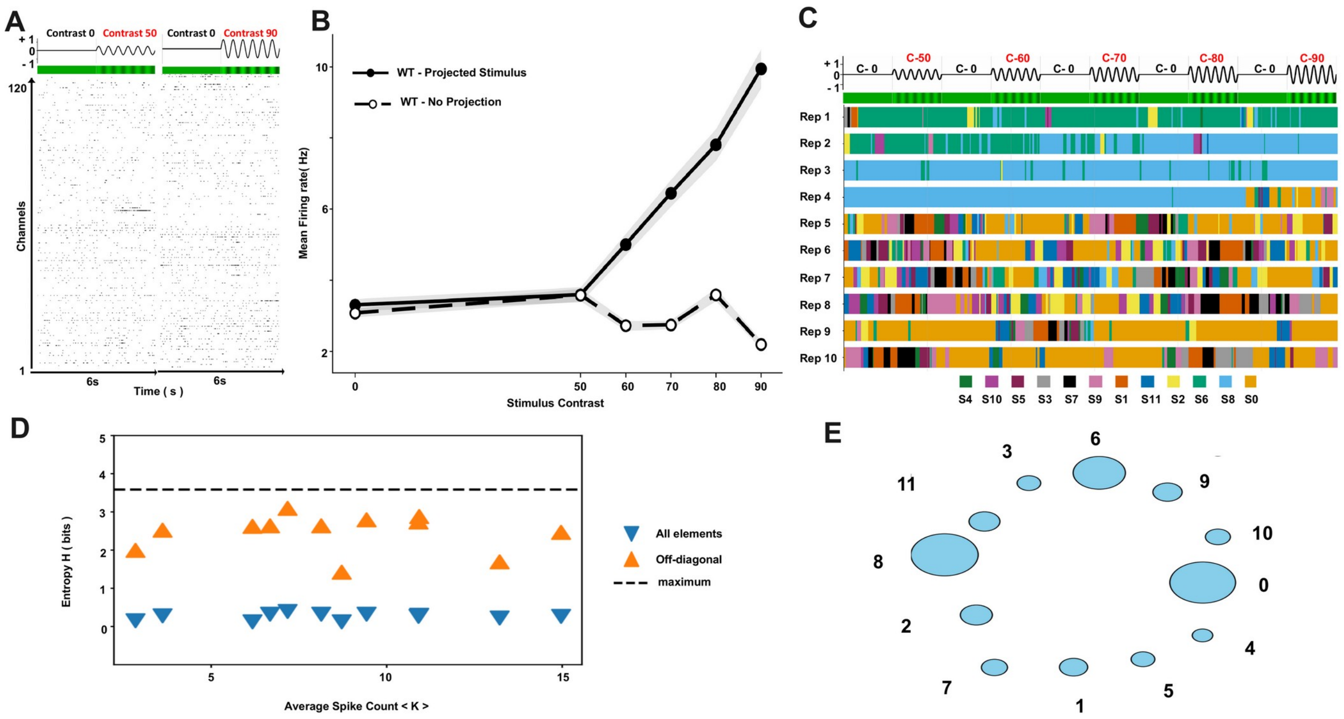


**Figure 5-2. No-stimulus control (recording WT3, no-projection)**: population dynamics with no visual projection onto the retina. (A) Population raster (120 channels) recorded across the same timing protocol used in the main dataset (nominal contrast blocks C-0 through C-90), but with no visual stimulus actually projected onto the retina, isolating any activity attributable to the recording/protocol timing itself rather than genuine light-evoked responses. (B) Mean firing rate (± SEM) plotted against the nominal contrast label of each timing block. Unlike the stimulus-driven data (Fig. 4B), firing rate does not increase monotonically with nominal contrast, indicating that the contrast-dependent recruitment seen in the main dataset depends on actual stimulus delivery rather than the timing structure of the protocol alone. (C) Viterbi-decoded state sequences (repetitions 1-10) of the same nominal timing protocol, colored by decoded state. State usage is markedly less diverse than in stimulus-driven recordings, with several repetitions dominated by a single state throughout, consistent with the absence of stimulus-locked population structure. (D) Transition-matrix entropy (bits) versus average spike count (⟨K⟩) for each state of the model fitted to this control dataset, shown separately for all transition-matrix elements (blue, downward triangles) and off-diagonal, state-switching elements only (orange, upward triangles), using the same convention as Fig. 5C. (E) Force-directed graph of the fitted state-transition structure for the control dataset, with node size proportional to state occupancy and edge width proportional to transition probability. The no-projection recording was made in the dark, with the same nominal stimulus-timing protocol applied to define analysis windows as in the stimulus-driven recordings, so the two conditions could be compared using identical windowing. The error band in (B) is SEM; off-diagonal probabilities in (D) are renormalized, and the edge-display threshold in (E) is 0.05, matching Fig. 2C. No off-diagonal edge exceeds this display threshold, which does not imply zero transition probability.

#### Figure 5-3

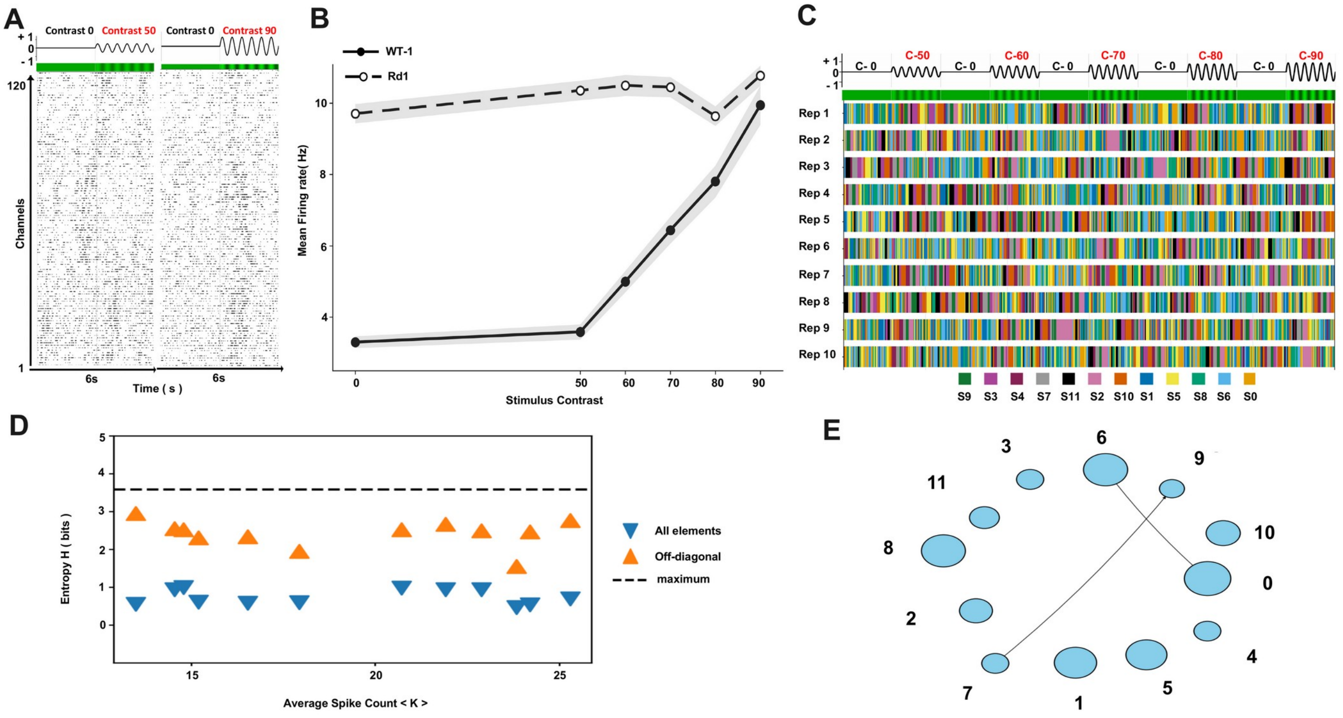


**Figure 5-3.** PHASOR outputs for the rd1 recording (rd1-1; C3H/HeJ, male, left retina, P105): population dynamics in a photoreceptor-degenerate retina, recorded with the stimulus projected exactly as in the wild-type recordings. This complements the no-projection recording in Figure 5-2: there the stimulus was withheld from an intact retina; here it was delivered to a degenerate retina. This single comparison does not separate the rd1 mutation from strain background or other differences between animals. (A) Population raster (120 channels) across the full stimulus protocol (contrast blocks C-0 through C-90, f0 = 2 Hz). (B) Mean firing rate (± SEM) as a function of stimulus contrast for the rd1 recording (open circles; n = 255 units), plotted against a representative wild-type recording (WT1, run 1; filled circles; n = 276 units) recorded under the identical protocol. Wild-type firing rate rises systematically with contrast; rd1 firing rate is already elevated at 0% contrast and remains largely contrast-invariant, showing no comparable stimulus-driven recruitment. (C) Viterbi-decoded state sequences for the rd1 recording across repetitions 1-10 of the full stimulus protocol, colored by decoded state (12-state model, S0-S11). All states are used, but the sequences do not realign at contrast-block boundaries, consistent with the absence of stimulus-locked population structure. (D) Transition-matrix entropy (bits) versus average spike count (⟨K⟩) for each state of the model fitted to this recording, shown separately for all transition-matrix elements (blue, downward triangles) and off-diagonal, state-switching elements only (orange, upward triangles), using the same convention as Fig. 5C. (E) Force-directed graph of the fitted state-transition structure, with node size proportional to state occupancy and edge width proportional to transition probability. The error band in (B) is SEM across units within this single recording and is not a measure of biological replication; off-diagonal probabilities in (D) are renormalized, and the edge-display threshold in (E) is 0.05, matching Fig. 2C. Recording metadata and fitted hyperparameters are listed in Table 2.
